# Neural sequences in primate prefrontal cortex encode working memory in naturalistic environments

**DOI:** 10.1101/2022.08.18.504406

**Authors:** Megan Roussy, Alexandra Busch, Rogelio Luna, Matthew L. Leavitt, Maryam H. Mofrad, Roberto A. Gulli, Benjamin Corrigan, Ján Mináč, Adam J. Sachs, Lena Palaniyappan, Lyle Muller, Julio C. Martinez-Trujillo

**Author notes:** Co-first author (Megan Roussy, Alexandra Busch). Co-senior author (Lyle Muller, Julio C. Martinez-Trujillo). MR, LP, and JCMT planned the experiments. MR and RL trained animals, performed the experiments and conducted data preprocessing including spike sorting. MR and AB analyzed the data, created figures and wrote the paper with help from JCMT and LM. BC developed unique MATLAB code for eye movement classification and analysis. RAG developed code for data preprocessing and contributed essential knowledge of experimental design and data analysis. BC and RAG trained animals to perform eye fixations for eye tracking calibration. MLL collected the ODR dataset. AJS, JCMT, RAG, RL, and MR planned and conducted surgeries. JM contributed to the dimensionality analysis development. MHM contributed to data analysis and analysis planning.

## Abstract

Working memory is the ability to briefly remember and manipulate information after it becomes unavailable to the senses. The mechanisms supporting working memory coding in the primate brain remain controversial. Here we demonstrate that microcircuits in layers 2/3 of the primate lateral prefrontal cortex dynamically represent memory content in a naturalistic task through sequential activation of single neurons. We simultaneously recorded the activity of hundreds of neurons in the lateral prefrontal cortex of macaque monkeys during a naturalistic visuospatial working memory task set in a virtual environment. We found that the sequential activation of single neurons encoded trajectories to target locations held in working memory. Neural sequences were not a mere successive activation of cells with memory fields at specific spatial locations, but an abstract representation of the subject’s trajectory to the target. Neural sequences were less correlated to target trajectories during perception and were not found during working memory tasks lacking the spatiotemporal structure of the naturalistic task. Finally, ketamine administration distorted neural sequences, selectively decreasing working memory performance. Our results indicate that neurons in the lateral prefrontal cortex causally encode working memory in naturalistic conditions via complex and temporally precise activation patterns.

## Introduction

Working memory is the ability to briefly maintain and manipulate information ‘in mind’ to achieve a current goal (Baddeley, 1986; Baddeley, 2003). Brain circuits supporting working memory differ from those for sensory processing in that they must represent precise information in naturalistic contexts in the absence of sensory inputs (Goldman-Rakic, 1990; Arnsten, Wang, & Paspalas, 2012; Wang, 2021; Roussy et al., 2021a). They also differ from long-term memory circuits in that the information is only maintained for short time intervals-just long enough to complete a specific task. Despite five decades of study, the neural mechanisms underlying working memory remain controversial.

The primate lateral prefrontal cortex (LPFC) has been widely implicated in working memory function as evidenced by previous lesion and electrophysiological studies in macaque monkeys (Curtis & D’esposito, 2004; Constantinidis., et al., 2018; Leavitt et al., 2017a; Pasternak et al., 2015). A long-supported mechanism for coding of working memory representations in LPFC of primates during delayed response tasks is persistent firing in single neurons selective for the memorized information (Fuster & Alexander, 1971; Constantinidis., et al., 2018). During such tasks, subjects must remember the location or features of a sample item for a few seconds after its disappearance, and then produce a behavioral response, e.g., a saccade to a remembered location. However, most delayed response tasks used to explore the neural mechanisms of working memory lack the spatiotemporal structure of naturalistic behavior (i.e., they use simple stationary displays and constrain eye movements during memory maintenance). During many natural behaviors involving working memory eye gaze is unconstrained and the visual scenery is rich and dynamic (Roussy, et al., 2021a).

Studies using delayed response tasks with increased spatiotemporal complexity report few single neurons with persistent firing during the entire delay period. Instead, many neurons fire transiently, during brief time intervals (Batuev et al., 1979; Lundqvist et al., 2016; Roussy et al., 2022). Thus, researchers have proposed alternative mechanisms to persistent firing, such as short-term synaptic storage (Stokes, 2015; Pals et al., 2020), or dynamic coding (Lundqvist et al., 2016; Parthasarathy et al., 2019). However, evidence in favor of such mechanisms is highly debated (Wang, 2021). Here, we hypothesize that a mechanism for working memory coding in naturalistic conditions must preserve the spatiotemporal structure of natural behavior while being robust to interference by concomitant sensory and motor signals. We specifically hypothesize that working memory coding during naturalistic tasks, in the presence of eye movements and rich visual scenery, relies on sequential activation of neurons in primate LPFC.

Neuronal sequences, consisting of temporally precise patterns of neural activity, have been reported to encode the varying spatiotemporal structure of motor signals in the high vocal center (HVC) of songbirds (Chi & Margoliash, 2001; Tang et al., 2014; Srivastava et al., 2017; Okubo et al., 2015; Daliparthi et al., 2019), and of spatial trajectories to remembered locations during navigation in the parietal cortex (Harvey et al., 2012) and the hippocampus of rodents (Itskov, et al., 2011; Eichenbaum, 2014; Zhou et al., 2020). Early investigations in macaque monkeys suggested that the spiking activity of a few single neurons in LPFC could have a precise and informative spatiotemporal structure (Abeles et al., 1993). However, sequences of single unit spiking activity have not been directly observed or causally linked to working memory during naturalistic behavior in primates (Wang, 2021).

We use microelectrode arrays to record neuronal activity in LPFC layers 2/3 of macaque monkeys during a naturalistic working memory task set in a 3D virtual environment. We find that temporally precise sequential patterns of neural activity in LPFC, extending over behaviorally relevant timescales of several seconds, represent important task variables for the successful maintenance of and navigation to remembered target locations in the 3D environment. These neural sequences robustly and flexibly represent trajectories to remembered locations during shifts in eye positions toward various elements of the environment. Sequences were not found when we examined an oculomotor delayed response task (ODR) which has been commonly used to explore working memory in previous studies. Further, pharmacological blockade of NMDA receptors with sub-anesthetic doses of ketamine demonstrates a causal link between sequences and working memory.

## Results

We trained two rhesus macaque monkeys on a visuospatial working memory task that took place in a virtual circular arena containing naturalistic elements (see Fig. 1a, b). We recorded neuronal activity using two 96-channel microelectrode Utah Arrays (Blackrock Neurotech, UT, USA) implanted in the left LPFC of both animals (Brodmann area 8a, 9/46 (Petrides, 2005)) (see Fig. 1c). The task began with a three second presentation of a target in one of nine possible locations in the arena (cue epoch). The target then disappeared, and after a two second delay period, the animal was required to navigate towards the cued target location using a joystick (see Fig. 1d). Virtual navigation within the environment was exclusively available during the navigation epoch. Animals were able to successfully perform this naturalistic working memory task (average correct trial rates across sessions were: NHP B: mean = 87%, NHP T: mean = 57%; chance = 11%) (Fig. 1e; Extended Data Fig. 1a-d). Eye movement was recorded throughout the task using a video eye tracker. Animals made frequent saccades to explore different scene elements throughout all trial epochs. Firing rates across the recorded neuronal population were poorly tuned for the direction and amplitude of saccades (Roussy et al., 2021b; Roussy et al., 2022) (see Extended Data Fig. 1e-k for eye behavior analyses).

**Fig. 1.**
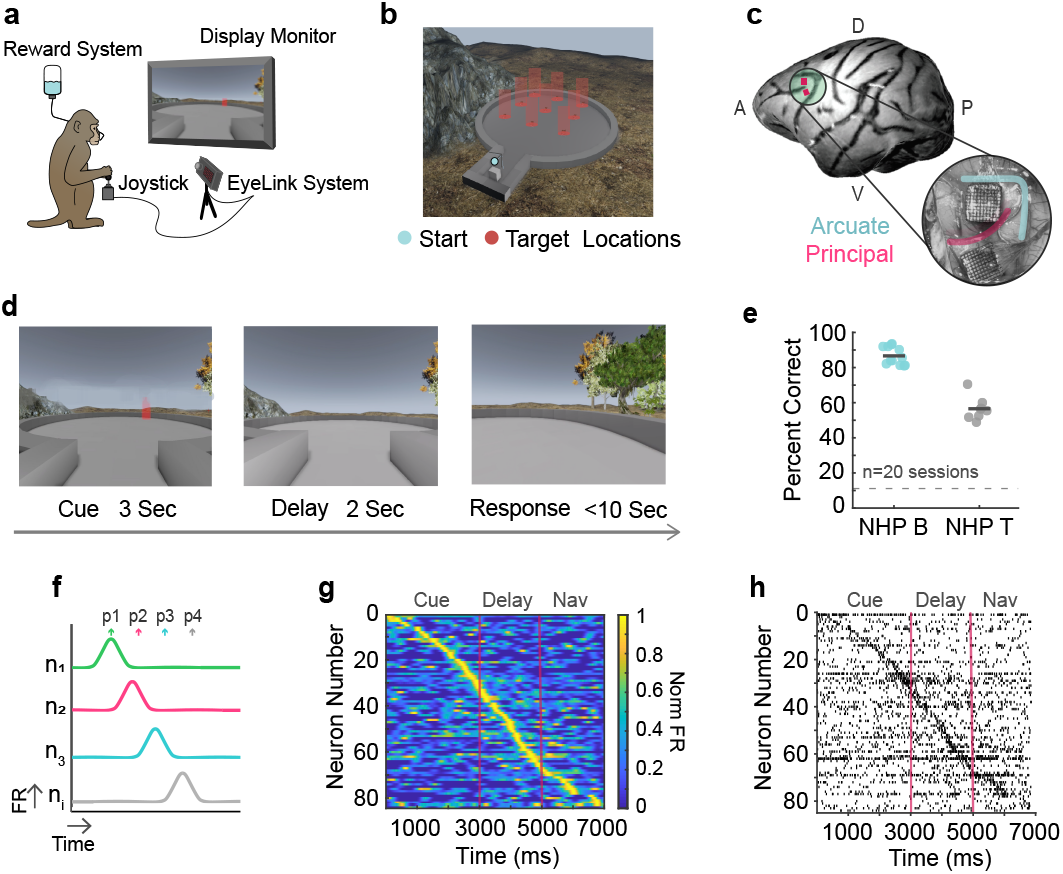
Experimental design. **a**, A monkey depicted in the virtual reality experimental setup. **b**, Overhead view of the nine target locations in the virtual environment. **c**, Locational representation and surgical image of the two Utah arrays implanted in the left LPFC of NHP T. **d**, Working memory trial timeline. **e**, Percent of correct trials for NHP B and NHP T. The dark gray lines represent mean values per animal and the gray dashed line represents chance behavioral performance. Data points represent data from individual sessions. **f**, Illustration of temporally tiled activation of individual neurons which may generate sequential patterns of activity at the population level. **g**, Normalized firing rates for simultaneously recorded neurons over trial time in one trial. **h**, Raster plot for the same example trial as ‘g’ in which each small vertical line represents an action potential. Pink vertical lines separate the task epochs.

### Neural sequences in LPFC neurons

Precise patterns of neural activity have been identified as a mechanism for representing complex processes in mammalian brains (Buzsáki, 2010); however, such patterns have not been identified during visuospatial WM tasks in primates. We hypothesize that WM representations during our naturalistic task are maintained by temporally precise neural sequences (Fig. 1f). Neural sequences are typically described by temporally precise activation of neurons above their background rates of activity. We observed that LPFC neurons exhibited brief (duration of 80% of max firing value = 220 ms) elevations of spike rate above their background levels of firing (Extended Data Fig. 2) at specific points during the task. To identify potentially relevant population-level patterns in these elevations of spike rate, we sorted neurons by their normalized peak firing time. Sequential patterns emerge in single trials, visualized here using spike density functions (Fig. 1g) and population rasters (Fig. 1h).

A code that relies on neural sequences implies temporally precise activation of single neurons (see Fig. 2a for schematic) (Buzsáki, 2010; van der Meij & Voytek, 2018). We examined the firing properties of 3543 neurons in 17 recording sessions (mean of 208, median of 229 simultaneously recorded neurons per session). Many neurons transiently fired during the same time in single trials of the same target condition (Fig. 2b, c, d, more examples in Extended Data Fig. 3). To quantify this regularity, we calculated the standard deviation (time consistency) of peak firing time between correct trials of the same condition for each neuron (Extended Data Fig. 4a). 20% of neurons (699 neurons) demonstrated a standard deviation below 1000 ms and 65% (2297 neurons) demonstrated a standard deviation below 1500 ms.

**Fig. 2.**
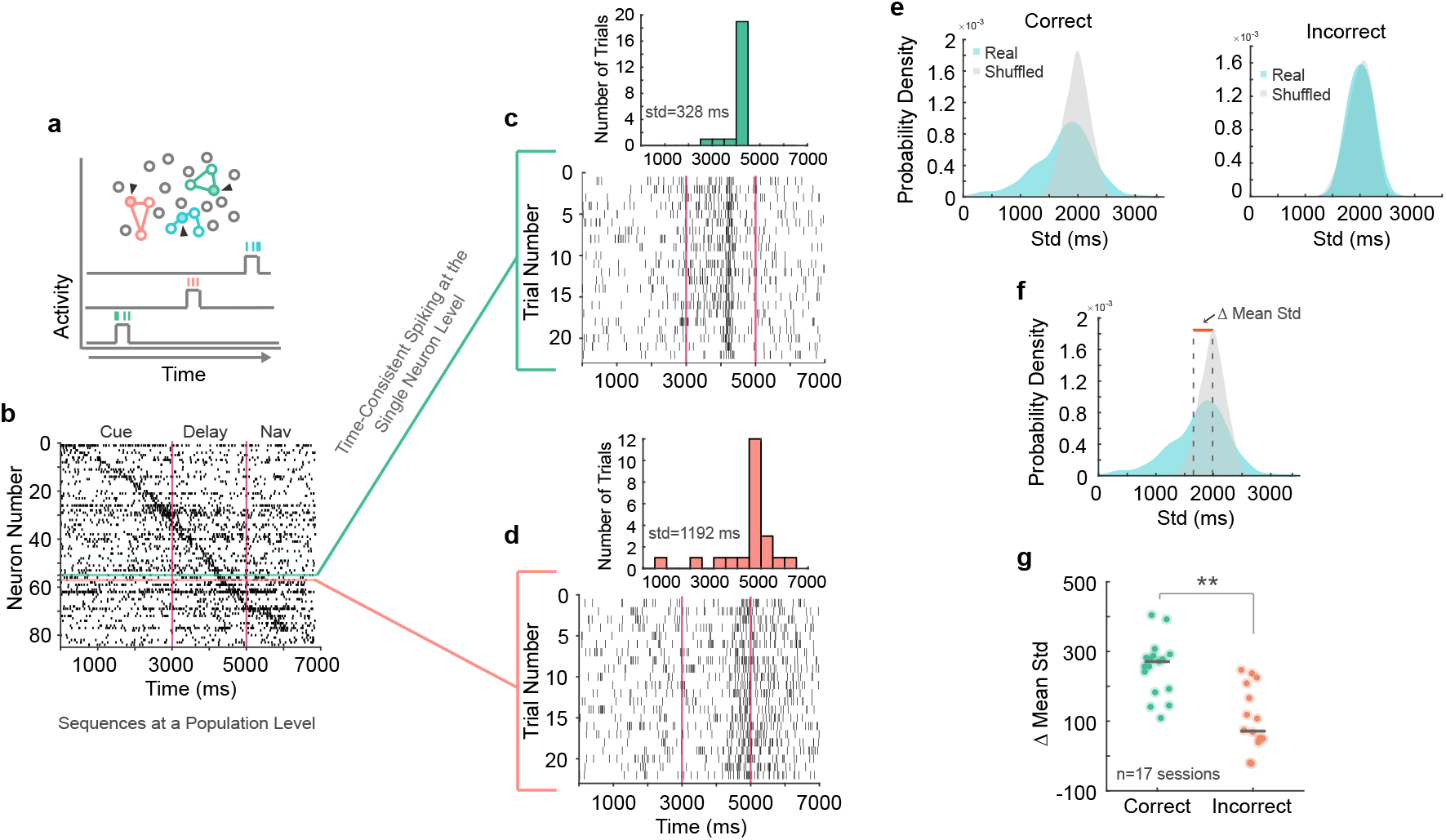
Time consistent neurons underlie sequence formation. **a**, Depiction of neural ensembles that are activated at different time points throughout a trial. Activity of a single neuron within the ensemble is represented by an increase in activity at a precise time point. Black arrows represent recording electrodes. **b**, Single trial raster showing the activity of all simultaneously recorded neurons. The green and pink horizontal lines highlight two example neurons. **c**, Example neuron one. The raster plot displays action potentials over trial time for this neuron over all trials in a certain condition. The inset histogram shows the number of trials in which the max firing time falls within a certain trial time. **d**, Represents the same information as ‘c’ for a second example neuron. **e**, Real and shuffled distributions of correct and incorrect trial-trial standard deviations of max firing time for all neurons in an example session. **f**, same as ‘e’ for correct trials. Dashed gray lines represent distribution means and the orange line indicates the difference in distribution means. **g**, Difference in real and shuffled distribution means for correct and incorrect trials. Dark gray lines represent median values per group and each dot represents data from a different session. *p<0.05, **p<0.01, ***p<0.001.

We additionally shuffled the peak firing times for each neuron within each trial to generate random firing time estimates across trials. The distributions of standard deviations for correct trials were shifted to lower values relative to the corresponding shuffled distributions (example session in Fig. 2e, all neurons in Extended Data Fig. 4a). The area of non-overlap between the lower tails of the real and shuffled distributions represents the neurons with peak firing times occurring more regularly than expected by chance within trials of the same condition. On the other hand, the real and shuffled distributions overlapped considerably for incorrect trials (example session in Fig. 2e; all neurons in Extended Data Fig. 4a), suggesting that neurons’ peak firing occurred at less consistent times during single trials of the same condition when animals made mistakes. Indeed, the difference between means of the real and shuffled distributions (Fig. 2f) was lower for incorrect trials than correct trials (correct: median = 270.9 ms, incorrect: median = 71.4 ms. Wilcoxon Signed-Rank Test, *p* = 0.001) (Fig. 2g; Extended Data Fig. 4b, c).

Using 11 sessions in which there are sufficient correct and incorrect trials in all nine conditions, we show that the standard deviation of neurons’ peak firing time (n = 2051) during correct trials (mean = 1358 ms) is significantly lower than incorrect trials (mean = 1828 ms; 1-way ANOVA, post-hoc, *p* = 3.8E-09; Extended Data Fig. 4a). This suggests that increased temporal precision of firing in single neurons is needed for correct task performance.

### Neural sequences are predictive of trajectory to remembered targets

We have demonstrated a pattern of sequential activity that spans the trial duration and is driven by the temporal consistency of the firing neurons. Next, we examined whether these identified sequences are related to working memory; more precisely, whether sequences can encode the contents of working memory during the delay epoch of the task, when the cue has disappeared, and navigation was not permitted. We developed a computational method to analyze spike sequences in single trials, allowing for efficient unsupervised discovery of neural sequences that are consistent within the same target condition. We represented individual sequences of peak firing during the delay epoch in each trial across the population of recorded neurons as complex-valued vectors. We performed dimensionality reduction on the resulting correlation matrix (Fig. 3a; Extended Data Fig. 5a-c). The resulting component values are projected into a 3-dimensional space where each colored circle represents a cluster centroid for a different target condition (Fig. 3b).

**Fig. 3.**
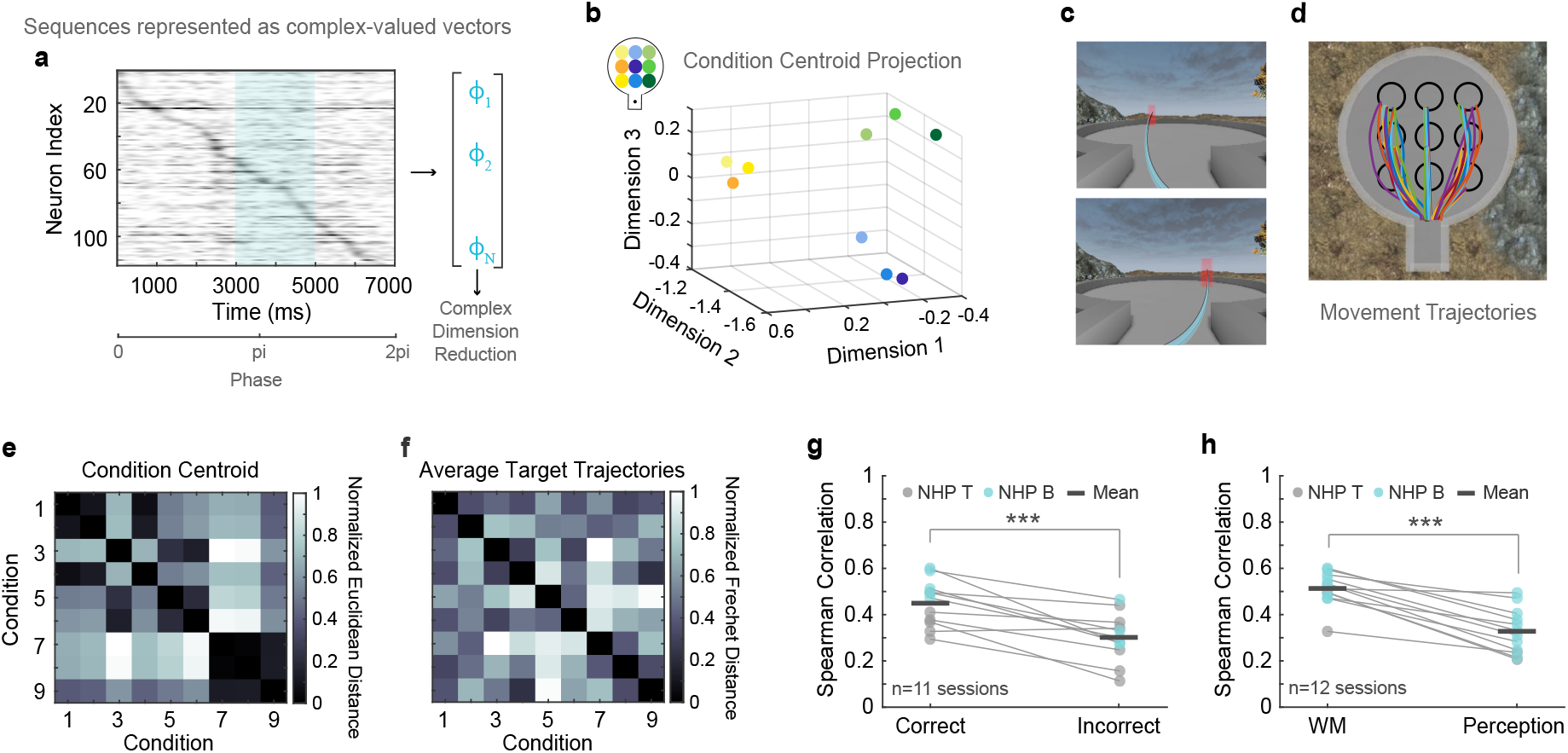
Neural sequences represent working memory content. **a**, Representation of max firing times per neuron during delay being converted to complex phase values and used in our complex vector dimensionality reduction method. **b**, Condition cluster centroids projected in 3D space. Centroid colors correspond with their position in the virtual environment (see inlet). **c**, Depiction of trajectories to two target locations. **d**, Example trajectories to the 9 target locations. Each line represents a trial. **e**, Coefficient matrix of Euclidean distances between condition centroids. **f**, Coefficient matrix of Frechet distances between average target trajectories for each target location. **g**, Correlation values for all sessions for correct and incorrect trials colored by animal. Dots represent individual sessions and matching sessions are connected by lines. **h**, Correlation values for all working memory and perception sessions colored by animal. Dots represent individual sessions and matching sessions are connected by lines. *p<0.05, **p<0.01, ***p<0.001.

Using a supervised distance-based classifier, we could correctly predict target condition within a single trial based on the clustering of condition centroids during the delay epoch (9 locations: median = 15% above chance, *p* = 9.7E-05; Extended Data Fig. 5d, see Methods - Projection Classification Analysis). We then observed that the condition centroids reliably formed three distinct groups based on the three primary trajectory directions to targets (i.e., left, center, right). Based on this observation, we hypothesized that this grouping may relate to task behavior since movement trajectories typically fall within these three directions. A supervised classifier based on this hypothesis can correctly predict target condition column (left, center, right) based on delay-epoch spiking activity within a single trial (median = 40% above chance, *p* = 1.7E-05; Extended Data Fig. 5e). Further, an unsupervised classifier developed from our analysis could predict the target column within a single trial based on the emergent clustering of projected data into column-based clusters - without any training required (median = 33% above chance, *p* = 1.7E-05; Extended Data Fig. 5f). Taken together, these results demonstrate that these patterns of spiking activity contain a unique temporal structure for different target conditions that may be related to remembered target locations.

To explore the direct relationship between sequences and task relevant behavior during working memory, we compared the distances between centroids during the memory delay epoch to distances between movement trajectories (i.e., the trajectories the animals used to reach the remembered target location). We calculated the Spearman correlation between matrices containing the Euclidean distance between condition centroids, and the Frechet distance between average traveled trajectories to targets in the virtual arena (see Fig. 3c-f; see Extended Data Fig. 6 for alternative methods). The Frechet distance between two trajectories is a measure of similarity between them that takes into account the location and ordering of the points along the trajectories (Alt & Godau, 1995). The distance matrices were more positively correlated compared to those obtained when shuffling the target locations, suggesting that the separation between neural sequences in multidimensional space parallels the discriminability between trajectories to targets held in working memory (observed: median = 0.50, shuffle: median = 0.34, Wilcoxon Signed-Rank Test: *p* = 0.02). Moreover, the relationship between sequences and target trajectories predicts whether information is successfully maintained during the working memory delay period, with higher correlations for correct than incorrect trials (correct: mean = 0.45, incorrect: mean = 0.30. T-test, *p* = 5.7E-04) (Fig. 3g).

We compared the correlation results during the working memory delay period with those from a temporally equivalent period of a perceptual task, where the target did not disappear during the entire trial and therefore the animals did not need to represent the trajectory in working memory. The correlation was higher during the working memory delay epoch than during the perceptual task control delay epoch (Fig 3h; Spearman Correlation; working memory: mean = 0.51 perception: mean = 0.33, T-Test, *p* = 1.4e-05), indicating sequences were more correlated to behavior during working memory.

We further asked whether behaviorally relevant sequences were composed of neurons considered tuned for the remembered target location using conventional criteria (i.e., differences in integrated firing rates amongst locations). Sequences composed of tuned neurons were equally correlated with behavior than sequences composed of untuned neurons (1-way Anova, *p* = 0.94; Extended Data Fig. 6d). The latter indicates that classically considered untuned neurons can take part in the sequences.

Single neurons show trial-to-trial variability in their responses. One may ask whether sequences could be robust to this phenomenon, (i.e., not all the neurons active during trial n sequences for one remembered target location will be active in trial n+1, n+2, etc). Remarkably, the correlation between neural sequences and target trajectories during correct trials remains stable even after removing 70% of neurons from the population. 80 – 90% of neurons must be removed for this correlation to significantly change, at which point the correct trial correlation becomes equal to the incorrect trial correlation (Extended Data Fig. 6d, e). The latter indicates sequences are robust to ‘single trial absentee neurons’ and therefore to trial-to-trial response variability.

Furthermore, neural sequences during working memory were different from those that occur during the cue and navigation epochs (Fig 4 a-c) and they were most predictive of target condition (Spearman Correlation; cue: mean = 0.46, delay: mean = 0.60, navigation: mean = 0.51, all: mean = 0.49; 1-way Anova, *p* = 0.03) (Fig 4d). Sequences were also more correlated to target trajectories when we limit a neuron’s contribution to a sequence to one epoch (i.e., one neuron is only allowed to participate in a single epoch sequence – considering a single max firing time) compared to contributing to multiple epoch sequences (i.e., allows for multiple peak firing times) (Spearman Correlation; single: mean = 0.48, multiple: mean = 0.37; T-Test, *p* = 0.006) (Extended Data Fig. 6c). This latter result indicates sequences are most informative when different neurons contribute to different epochs, suggesting unique contributions of neurons in different trial periods.

**Fig. 4.**
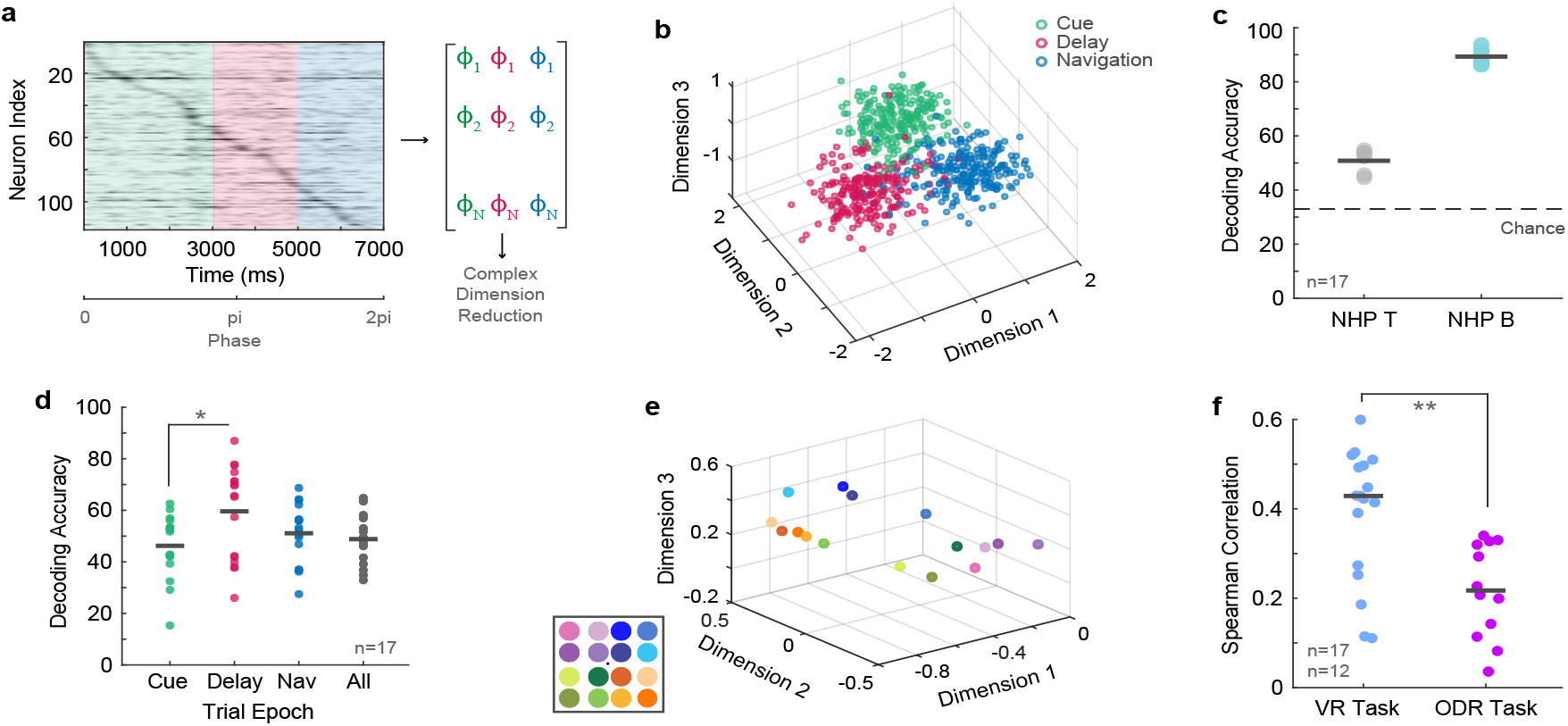
Working memory sequences are unique to naturalistic behavior. **a**, Representation of maximum firing times per neuron during the three task epochs being converted to complex phase values and used in our complex vector dimensionality reduction method. **b**, Epoch specific sequences projected in 3D space. Colored dots correspond to epochs and represent a single trial. **c**, Decoding accuracy using unsupervised classifiers for predicting the task epoch. Dots represent data from different sessions. **d**, Decoding accuracy using unsupervised classifiers for predicting the target column (left, middle, right) for sequences during task epochs. **e**, Condition cluster centroids for an ODR task projected in 3D space. Centroid colors correspond with their position in the virtual environment (see inlet). **f**, Correlation values for centroid distance and the distance between target locations for the virtual reality task and the ODR task. Dots represent data per session. *p<0.05, **p<0.01, ***p<0.001.

### Neural sequences are specific to naturalistic working memory

Neural sequences may operate as a mechanism for working memory coding in the naturalistic conditions of our task. To test this prediction, we conducted the same set of analyses exploring macaque LPFC single neuron temporal precision and population sequences in a classic oculomotor delayed response task (ODR) (Extended Data Fig. 7a, b). The task we used included 16 possible target locations. Here animals must fixate on a dot on a blank screen, then a peripheral target is flashed for a short time period. After target offset, the animals keep fixating on the dot for a few seconds while remembering the spatial location of the target. Upon the fixation dot offset, the animals make a saccade towards the remembered location to obtain a reward (Leavitt, 2017b, Leavitt, 2018). Saccades are ballistic movements that practically ‘teleport’ the fovea from the starting to the end point, without perception of the travelled path during eye movement (Bremmer et al. 2009).

As opposed to the VR navigation task, when neurons during the ODR task were ordered by peak firing time, the patterns of activation were often disrupted or incomplete (Extended Data Fig. 7c), suggesting that the organization of spiking activity may be different from the VR task. This may be related to neurons during the ODR delay epoch exhibiting less temporally consistent peak firing times from trial to trial. For many instances, real and shuffled distributions of standard deviations were overlapping (Extended Data Fig. 7d). Indeed, the difference in means between real and shuffled distributions was significantly smaller in the ODR task compared to our naturalistic VR task (ODR1: median = 93.2. ODR2: median = 31.6, VR: median = 270.9; Kruskal Wallis, *p* = 1.2e-06) (Extended Data Fig. 7e).

To further explore this issue, we applied the complex-valued dimensionality reduction analysis described above to the ODR task data. Condition centroids were clustered in quadrants based on position of target location as reported previously using spike rate-based analysis (Leavitt, 2018) (Fig. 4e). We calculated the correlation between the matrices of centroid distances and target locations Euclidean distances. The correlation was significantly smaller in the ODR than in the naturalistic VR task (ODR: median = 0.22, VR: median = 0.43. Wilcoxon Rank Sum, *p* = 0.004) (Fig. 4f).

These results indicate that sequences are more correlated to behavioral performance during the naturalistic VR tasks than during the classic ODR task used by previous studies. The naturalistic VR task is different in several ways. First, it measures visuospatial working memory in a dynamic and more spatiotemporally complex environment. Second, it allows for free visual exploration via saccades. Third, it requires 3D navigation to a target location. Neural sequences may be best utilized in the episodic and dynamic spatiotemporal context of our VR working memory task.

### Neuronal sequences represent abstract trajectoyr

The previous analyses demonstrate that working memory sequences contain information about the trajectories to remembered locations suggesting that sequences map into behavioral paths. Indeed, condition centroids were more highly correlated to the Frechet distance between the traveled trajectories to target locations (median = 0.50) than to the Euclidean distance between target locations (median = 0.43; Wilcoxon Sign Rank Test, *p* = 0.01); thus, sequences better represent trajectories to targets than target location alone. Real trajectories are also more correlated to condition centroids than ideal trajectories to targets (calculated by Euclidean distance from start to target location) (Extended Data Fig. 8a) (median = 0.43; Wilcoxon Sign Rank Test, *p* = 0.01). Here one must consider that traveled trajectories are imperfect and can be distinct from ideal trajectories. Real trajectories reflect idiosyncrasies of remembered trajectories and the virtual environment, and may reflect perceived curvature of the arena and obstacles in space (i.e., arena walls) (see Fig. 3d for example trajectories).

One may argue that the observed sequences represent activation of neurons with mnemonic ‘place fields’ similar to sequential activity of place cells in the hippocampus (Itskov et al., 2011; Eichenbaum, 2014; Zhou et al., 2020). Inconsistent with this idea, the sequences are differentiable between memory delay and navigation evidenced through classification analysis (mean decoding = 76%, median decoding = 87%, compared to chance (33%): T-Test, *p* = 9.2e-08) (Fig. 4b, c).

If sequences were primarily reflecting motor planning during the delay period or neural replay of planned trajectories during the response/navigation period, one may anticipate neural sequences during the delay and response epochs from the same trial to be highly correlated, and that this correlation would be higher than sequences from different trials. This was not the case. Delay and navigation epoch sequences were equally correlated between different trials as they were within the same trial (Extended Data Fig. 8b, c). These results indicate that neural sequences in macaque LPFC represent remembered trajectories to target locations, and that such representation is specific to the working memory delay period of the task. The latter makes the representation distinct from classical replay found in structures such as the hippocampus (Skaggs et al. 1996). LPFC neurons appear to represent more abstract qualities of target trajectory.

### Ketamine disrupts neuronal sequences and impairs working memory performance

In order to demonstrate a causal link between neuronal sequences and working memory we used ketamine, a N-methyl-D-aspartate (NMDA) receptor noncompetitive antagonist that induces selective working memory deficits in humans and animals (Frohlich & Van Horn, 2014; Roussy et al., 2021b, Wang et al., 2013). We injected subanesthetic doses of ketamine (0.25 mg/kg – 0.8 mg/kg) intramuscularly while animals performed the task (see experimental timeline in Fig. 5a, Roussy et al., 2021b). Ketamine drastically reduced performance of our virtual working memory task without affecting performance on a perception control task. Working memory performance recovered 30 minutes to 1 hour post-injection in the late post-injection period (Pre-Injection: median = 77%, Early Post-Injection: median = 28%, Late Post-Injection: median = 66%; Kruskal Wallis, *p* = 8.5e-05) (Fig. 5b; Extended Data Fig. 9a, b).

**Fig. 5.**
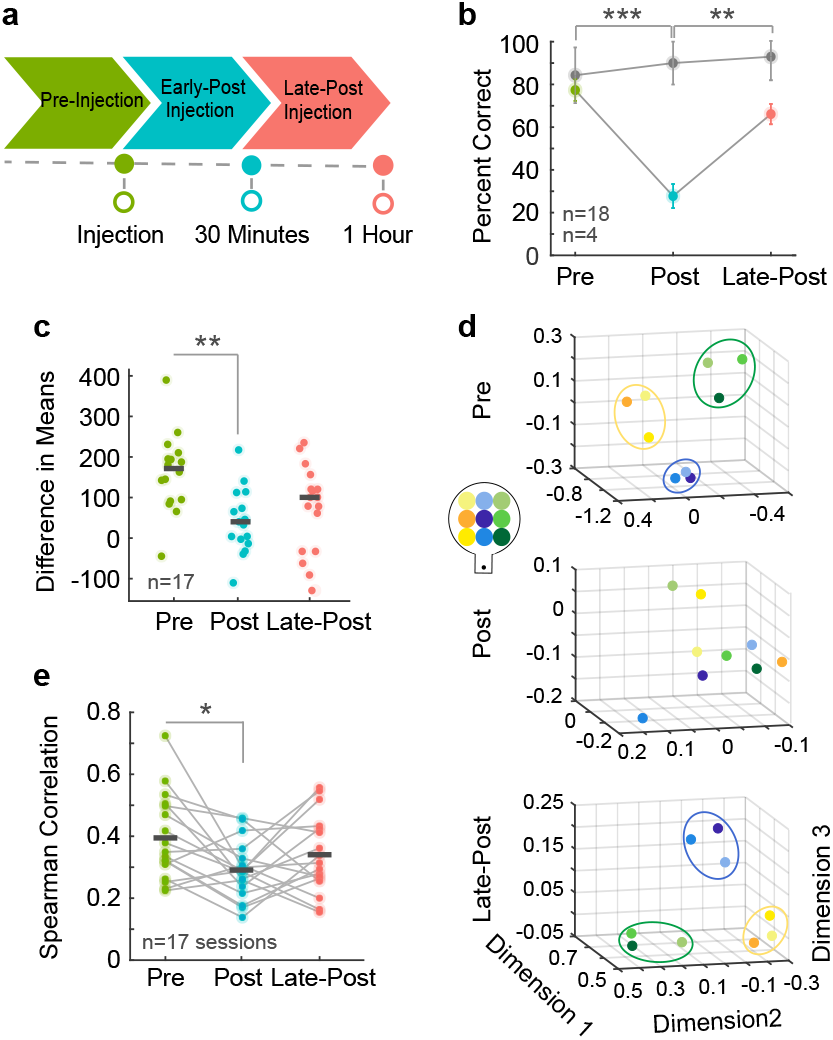
Ketamine manipulation distorts neural sequences and working memory. **a**, Experimental timeline for ketamine injection. Pre-injection period is depicted in green, early-post injection in blue, and late-post injection (recovery) in pink. **b**, Task performance as percent of correct trials for each injection period. Color dots represent median values per injection period for working memory data and gray dots represent median values per injection period for perception control data. Asterisks indicate significance between working memory injection periods. **c**, Difference in real and shuffled distribution means between ketamine injection periods. **d**, Condition centroids projected in 3D space. Centroid colors correspond with their position in the virtual environment (see inlet). Ellipsoids are illustrated guides to indicate behaviorally relevant groupings of targets in the pre-injection and late post-injection periods. These groupings are notably absent in the post-injection period. **e**, Spearman correlation values for each ketamine injection period. Dots represent individual sessions and matching sessions are connected by lines. *p<0.05, **p<0.01, ***p<0.001.

After ketamine injection, the differences in standard deviation distribution means for peak firing times between the real and shuffled data decreased suggesting that neurons fired with less time consistency after ketamine (Pre-Injection: median = 171.6, Early Post-Injection: median = 40.2, Late PostInjection: median = 100.4; Kruskal Wallis, *p* = 0.002) (Fig. 5c; Extended Data Fig. 9c, d). Behaviorally relevant groupings of condition centroids were similar between the non-injection data set and the pre-injection ketamine data set (Fig. 3b; Fig. 5d). This grouping was lost after ketamine injection but was regained 1 hour later as behavioral performance recovered (Fig. 5d). We also saw that the correlation between condition centroid distances and target trajectory distances decreased after ketamine and then recovered, indicating that sequences were less predictive of remembered target location immediately after ketamine injection (Pre-Injection: mean = 0.39, Early Post-Injection: mean = 0.29, Late Post-injection: mean = 0.34. 1-way Anova, post-hoc, *p* = 0.04) (Fig. 5e; Extended Data Fig. 9e, f). There was no change in any of the described measures in a saline control condition (Extended Data Fig. 9g, h). These results indicate a causal link between NMDA receptor dysfunction caused by ketamine and disruption of neuronal sequences leading to deficits in working memory.

## Discussion

We recorded the responses of hundreds of single neurons in the macaque LPFC during a complex visuospatial working memory task set in a naturalistic virtual environment. We report three major findings: (1) sequences of population activity represented trajectories to remembered locations in the environment (2) neural sequences of single neuron spiking activity were predictive of behavioral performance (3) NMDA receptor antagonism by ketamine disrupted neuronal sequences, selectively impairing working memory performance.

### Neural sequences and working memory coding

Prefrontal neural activity during tasks that require holding a single item in working memory during a delay response period have demonstrated persistent activity that represents the memoranda (Leavitt et al., 2017a). One major shortcoming of the persistent firing hypothesis is that it may not be able to support working memory representations with rich spatiotemporal structure (Steveninck et al., 1997; Lestienne & Strehler, 1987; Lundqvist et al., 2016). Indeed, in tasks during which sequences of multiple items need to be held in working memory, persistent firing is rare (Lundqvist et al., 2016). A recent study reported that during a multi-item spatial working memory task in which monkeys had to remember a series of spatial locations in sequential order, temporally organized neuronal populations represented the order in which items were remembered (Xie et al., 2022). These studies demonstrate that additional mechanisms may be needed to support coding of working memory representations when the memoranda have spatiotemporal structure.

Our paradigm differs from those used in previous studies. We did not use multiple memoranda; instead, our subjects remembered a single target location and the trajectory to the location in a 3D virtual naturalistic environment. Importantly, our study did not restrain eye position, allowing for naturalistic exploration of the scene while information is being held in working memory. The rationale behind studies restraining eye position is to avoid the interference caused by eye position signals and changes in the retinal image and consequently in visual inputs, on the working memory representation (Suzuki & Gottlieb, 2013). However, in naturalistic conditions working memory coding must be robust to such changes. To our knowledge, working memory coding has not been tested under naturalistic conditions.

Previous studies in macaques have tried to approach the idea of transiently active neurons maintaining working memory through shared temporal relationships by exploring spike train patterns of several neurons. However, due to methodological constraints, these studies were unable to record large numbers of simultaneously active neurons and thus unable to demonstrate sequence coding (Prut et al., 1998). Our study has overcome this limitation by recording from hundreds of simultaneously active neurons, revealing precise sequences of single unit spiking activity that encode specific working memory content.

Studies in mice that simultaneously record from many neurons have reported neuronal activation sequences during short-term memory tasks in the posterior parietal cortex and dorsomedial striatum (Harvey et al., 2012; Akhlaghpour et al., 2016). In the rodent hippocampus, sequences of place cell activation signal trajectories to remembered locations that are stored in long-term memory (Skaggs & McNaughton, 1996). Thus, sequential activation of neurons to encode spatiotemporal episodes appears to be a general coding mechanism across species.

However, the neural sequences we report in this study differ in several ways from those described in the rodent. First, they occur in the LPFC, a brain area that appears during brain evolution in anthropoid primates (Passingham & Wise, 2012). More specifically, the sequences reported here occur within the supragranular layers 2 and 3, where working memory representations have been reported (Bastos et al., 2018; Finn et al., 2019). The expansion of layers 2/3 is found in anthropoid primates and is accompanied by changes in the morphology, size (Gilman et al. 2017) and proportion of different interneuron types (Torres-Gomez et al., 2019) relative to other species and brain areas. Thus, LPFC layers 2/3 may have evolved a microcircuitry for holding internal working memory representations that can be supported by persistent firing, when spatiotemporal structure is poor and concomitant visual and motor signals are not present; or by neuronal sequences, when spatiotemporal structure is rich and distracting signals are present, as in our naturalistic task.

We propose LPFC layers 2/3 neural sequences may allow primates to represent short-term spatiotemporal episodes ‘in the mind’. Such episodes can be dissociated from sensory and motor signals and may be key to an enriched virtual world that enables enhanced cognitive control, planning and creativity observed in anthropoid primates (Passingham & Wise, 2012). Importantly, this form of episodic working memory may correspond to the episodic working memory buffer proposed by theoretical and behavioral studies of working memory in humans (Baddeley, 2000).

### Causality and potential mechanisms

Through pharmaceutical manipulation, we identify that sequence generation relies on NMDA receptor function. The interactions between inhibitory interneurons and excitatory pyramidal cells plays an important role in LPFC prefrontal circuits during working memory tasks (Wang et al., 2004). Therefore, the precise activation of pyramidal cells may be dependent on a temporally coordinated ‘release of inhibition’ by interneurons (Cannon, et al., 2015; Kosche et al., 2015). We have demonstrated in past research that NMDA receptor antagonism using the same doses of ketamine as in this study selectively decreases the firing of narrow spiking neurons (Roussy et al., 2021b). A parsimonious explanation for our findings is that ketamine induced loss of firing in narrow spiking interneurons (e.g., PV basket or chandelier cells) which in turn impaired their ability to coordinate sequences in pyramidal cells ultimately causing deficits in working memory. The fact that the effect of ketamine was selective for the working memory task further support our view that the sequential activation mechanisms reported here is particularly important for mental representations that ‘live’ within the LPFC microcircuits.

There may be various benefits of a sequence-based code. It would be more energy efficient than one that relies on continuous activation of neurons in a population. A temporal code may also be robust to interference by other concomitantly occurring signals, as is the case during naturalistic tasks. Temporal specificity may also add complexity to prefrontal networks, allowing for higher dimensional representations and flexible cognition.

Finally, neuronal sequence codes in the LPFC could be the substrate of working memory episodes that can be played in the mind and the neural correlates of the episodic buffer component of working memory systems in the human brain (Baddeley, 2000). The episodic working memory buffer was proposed as an upgrade of the classic working memory model that contained a visuomotor sketchpad, a phonological loop and an executive or attentional controller (Baddeley, 1986; Baddeley, 2000). The episodic buffer can bind experiences into working memory episodes. Such episodes can exist temporarily ‘in the mind’ and be ‘erased’ without undergoing long term storage. The latter make them distinct from long term episodic memories (Tulving, 2002) which engage hippocampal circuits (Burgess, Maguire, & O’keefe, 2002).

### Conclusion

We demonstrate robust and behaviorally relevant temporal organization of spiking activity in layers 2/3 of LPFC during a naturalistic working memory task. Neuronal sequences during periods of working memory maintenance represent the spatiotemporal structure of the information held in working memory. Sequences were disrupted by low doses of ketamine which caused impaired behavioral performance. We conclude that layers 2/3 LPFC circuitry in primates contains the neural substrates for temporarily representing working memory episodes ‘in the mind’ without necessarily engaging sensory, motor and even long term memory systems. Such representations provide primates with a powerful tool for planning the future and adapting to the uncertainty of changing environments.

## Extended Data Figures

**Fig. 6.**
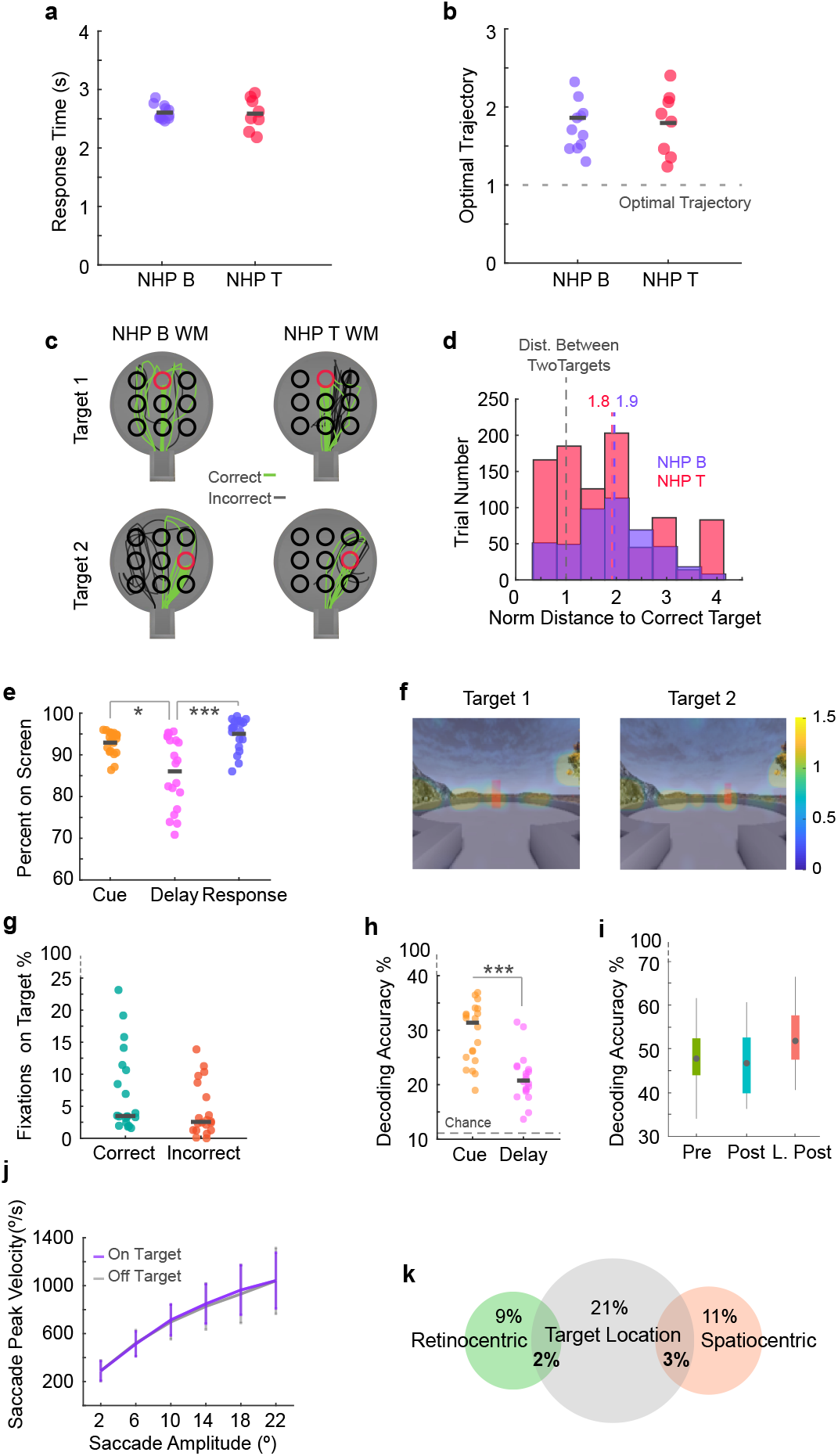
Task behavior. **a**, Response time for correct trials. Black lines indicate the median values for each animal. Dots represent data for each session. **b**, Optimal trajectory for correct trials. Calculated by real trajectory length/ Euclidean distance between the start location and target location. The dashed gray line indicates the optimal value of 1. Black lines represent median values and dots represent data for each session. **c**, Example animal trajectories for two target locations. Red circles indicate the target, green lines represent correct trials and gray lines represent incorrect trials. **d**, Distance from target for incorrect trials. Calculated as the Euclidean distance from the animal’s end location to the correct target location. Since ‘Unreal’ units are arbitrary, distance values are normalized by the distance (in ‘Unreal’ units) between two targets. A normalized distance of 1 indicated by the dashed gray line is the distance between two target centers. Dashed purple and red lines represent mean values. **e**, Percent of eye data points falling on screen for different task epochs. Black lines represent mean values for each group and dots represent data from each session. **f**, Heat maps of eye fixation position on screen during delay for two target examples. Eye fixation is concentrated in task relevant areas on screen but is not primarily concentrated on the target location. **g**, Percentage of total fixations during the delay epoch that land within the target location for correct and incorrect trials. The black lines represent median values and the dots represent data for each session. **h**, Decoding accuracy for predicting target location from eye fixation location during the cue and delay epochs. The black lines represent median values and the dots represent data for each session. The gray line indicates chance performance for 9 classes. **i**, Decoding accuracy for decoding target location from eye fixation position during the delay period for different ketamine injection periods. Decoding is shown for 3 classes (33.33% chance). The gray circles indicate median values. **j**, Main sequence values during the delay period for saccades that fall on and off target location. **k**, Proportion of neurons tuned for saccade landing position in retinocentric and spatiocentric reference frames as well as the proportion of neurons tuned for target location during the delay period. Overlapping sections indicate neurons that are selective for both remembered target location and saccade position. Panels ‘i’ and ‘k’ have been adapted from Roussy et al, 2021. Panels ‘a’, ‘b’, ‘e’, ‘f’, ‘g’, ‘h’, and ‘j’ have been adapted from Roussy et al., 2022 (bioArx). *p < 0.05, **p < 0.01, ***p < 0.001.

**Fig. 7.**
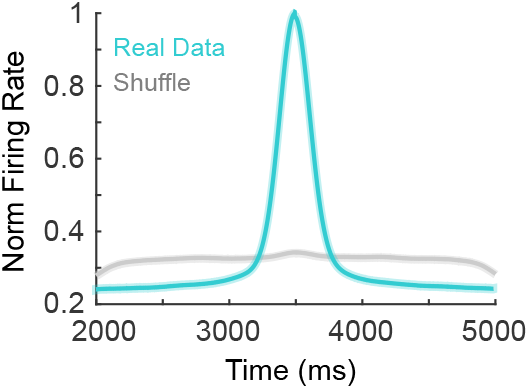
Neural sequences. Alignment of peak firing times for all neurons (blue) compared to shuffled peak firing times (gray).

**Fig. 8.**
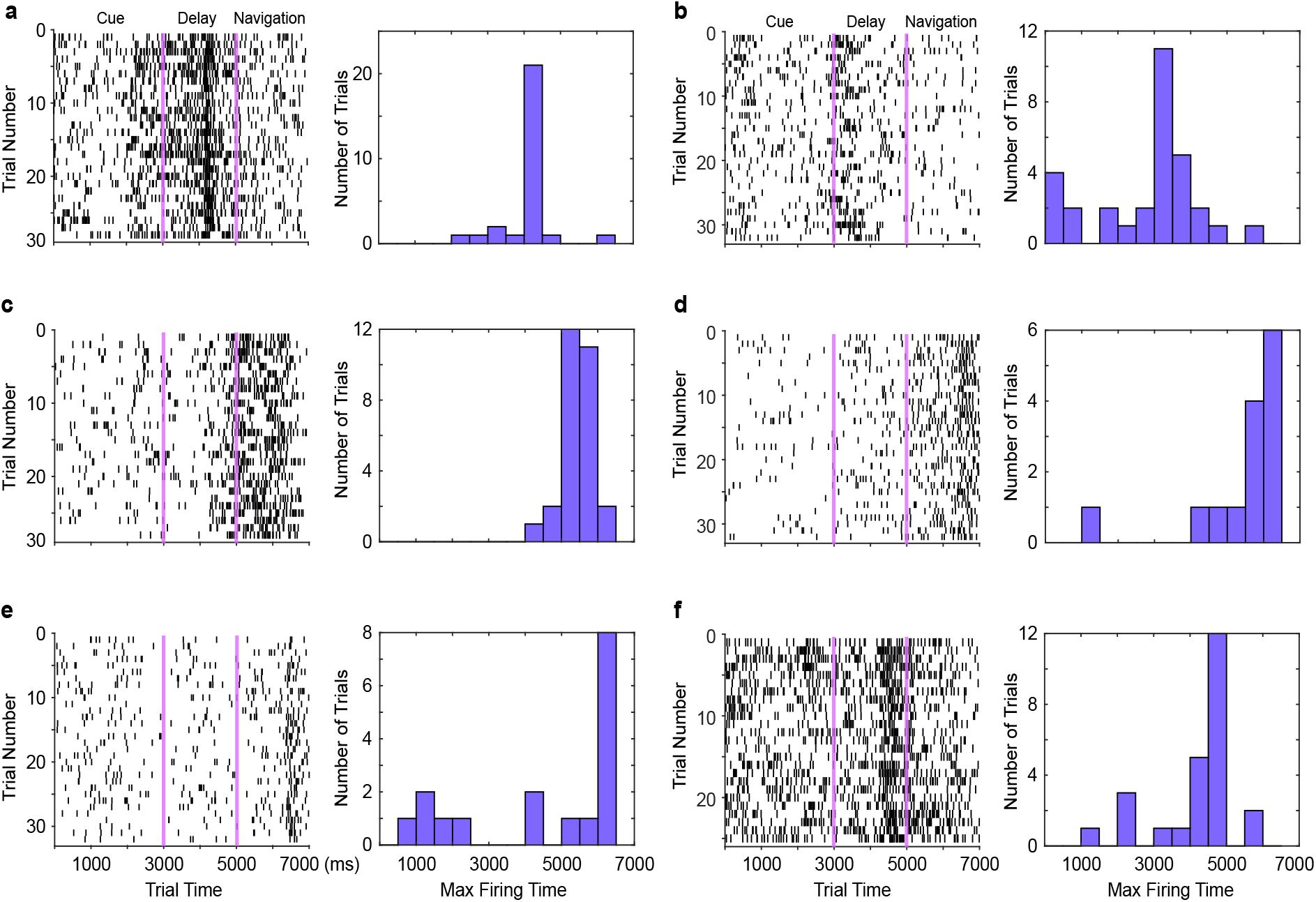
Example time consistent neurons. **a-f**, Example single neurons. Left column represents the activity of a neuron over trial time over all trials of a certain condition. Pink lines separate task epochs. Right column shows a histogram of max firing times per trial.

**Fig. 9.**
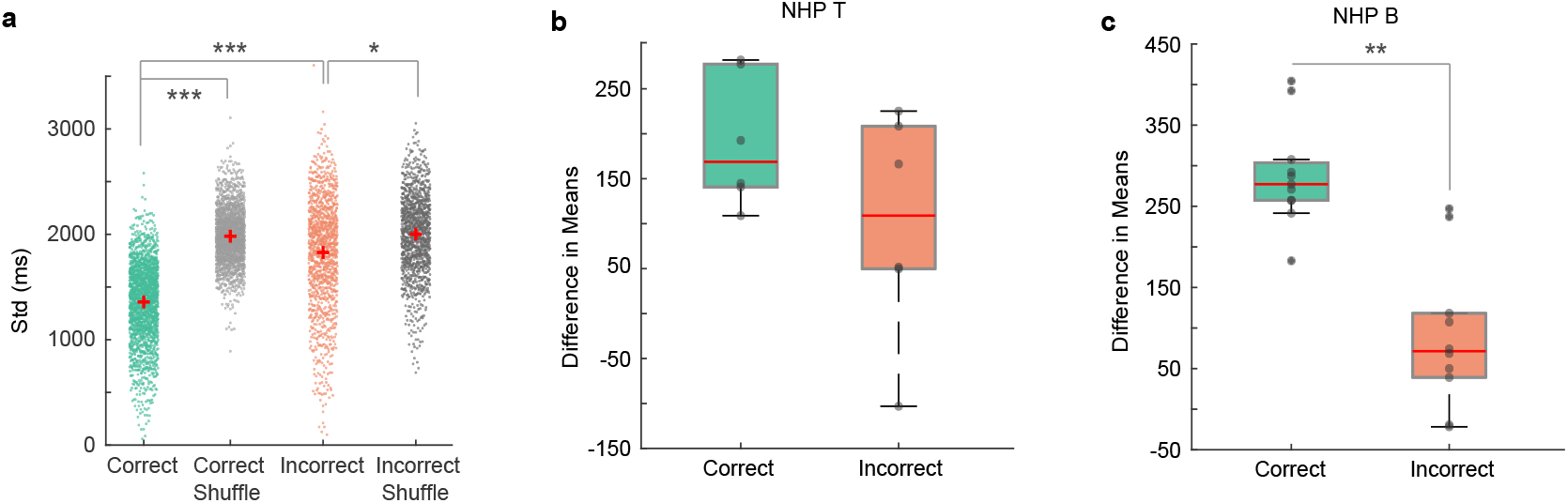
Time consistent neurons. **a**, Trial-trial standard deviation in max firing time for each neuron across sessions during correct and incorrect trials and for shuffled correct and incorrect data. The Red crosses indicate group means. **b**, Difference in real and shuffled distribution of the deviation in neuron action potential timing between trials. Presented for correct and incorrect trials for NHP T. The red lines represent median values. Dots represent data for individual sessions. **c**, Difference in real and shuffled distribution of the deviation in neuron action potential timing between trials. Presented for correct and incorrect trials for NHP B. The red lines represent median values. Dots represent data for individual sessions. *p < 0.05, **p < 0.01, ***p < 0.001.

**Fig. 10.**
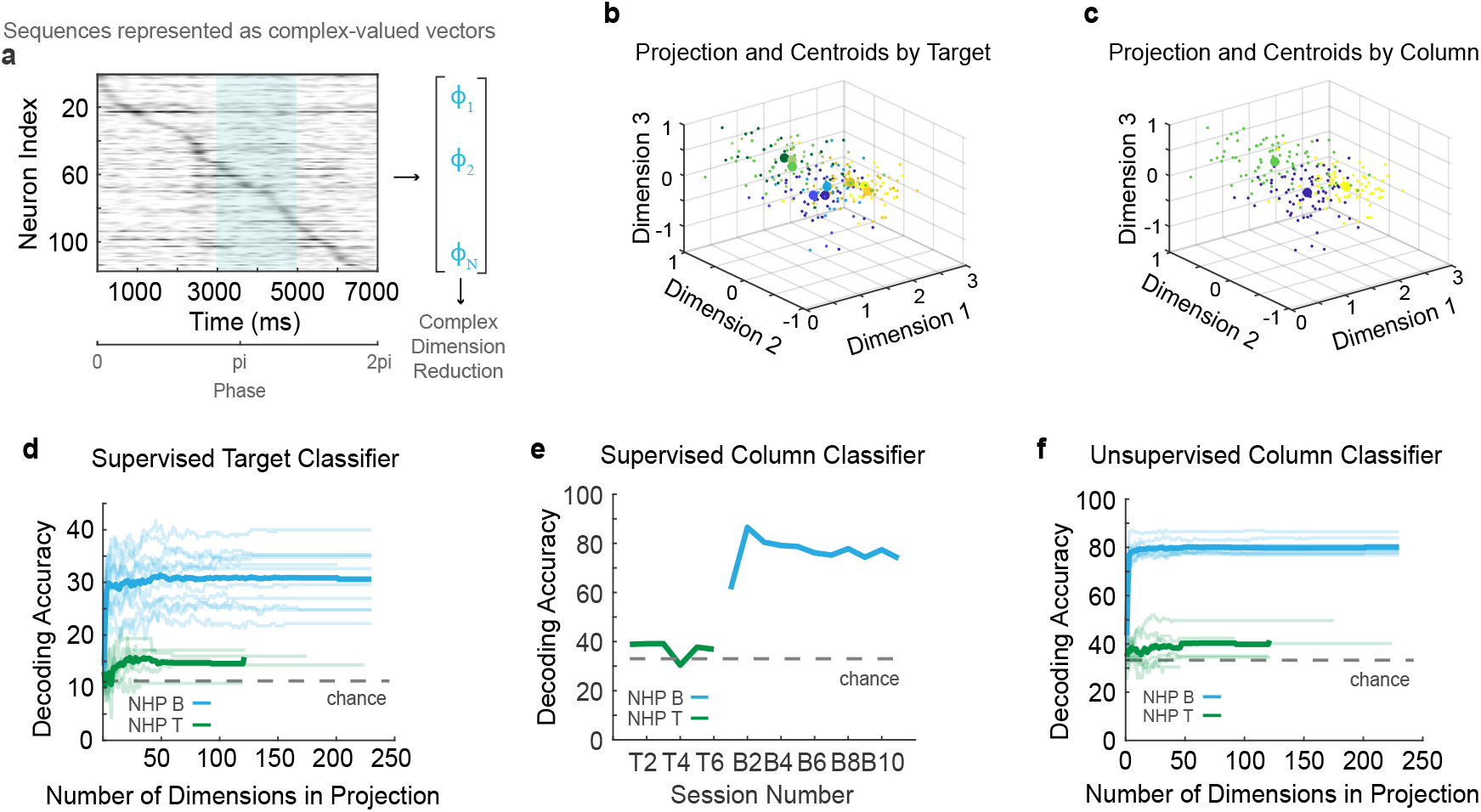
Classification of sequential coding. **a**, Representation of max firing times per neuron during delay being converted to complex phase values and used in our complex vector dimensionality reduction method. **b**, Large dots represent projected target centroids in 3D space. Smaller dots represent individual trials. **c**, Large dots represent projected column centroids in 3D space. Columns contain pooled trials between right, left, and center targets. Smaller dots represent individual trials. **d**, Supervised classification of target location using centroid distances. Each line represents decoding accuracy over number of dimensions considered for one session. **e**, Supervised classification of target column (left, right, center) using centroid distances projected into three dimensions. f, Unsupervised classification of target column (left, right center) using centroid distances. Each line represents decoding accuracy over number of dimensions considered for one session.

**Fig. 11.**
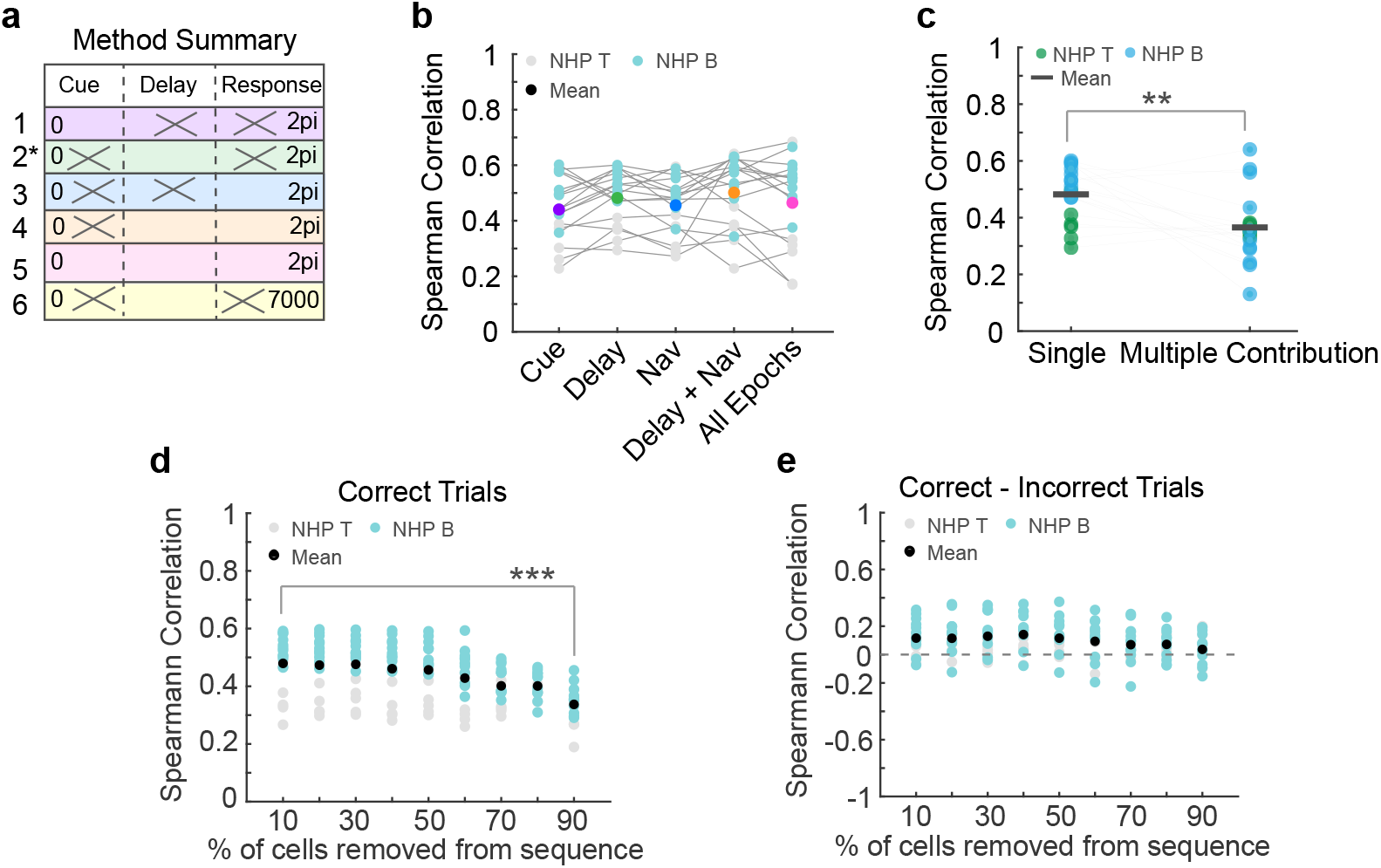
Deviations of correlation method. **a**, Method summary outlining different ways to calculate sequences. Crossed-out epochs indicate epoch data that was not used as part of the complex vectors for a given method. Asterisk indicates the method used in the main figures and text. **b**, Correlation based on each method outlined in ‘a’. Dots represent data per session. Colored dots represent mean and correspond to the table in ‘a’. **c**, Correlation when neurons are only considered in one epoch sequence or when all neurons participate in all sequences. This reflects the possibility of one instance of peak activity versus multiple occurrences of increased firing. Dots represent data per session. **d**, Spearman correlation using either tuned and matched untuned units. **e**, Spearman correlations for correct trials between delay neural sequences and target trajectories after removing 10 – 90% of neurons from the sequence. **f**, The difference between Spearman correlations for correct and incorrect trials after removing 10 – 90% of neurons from delay sequences. The dashed gray line represents 0 difference. *p < 0.05, **p < 0.01, ***p < 0.001.

**Fig. 12.**
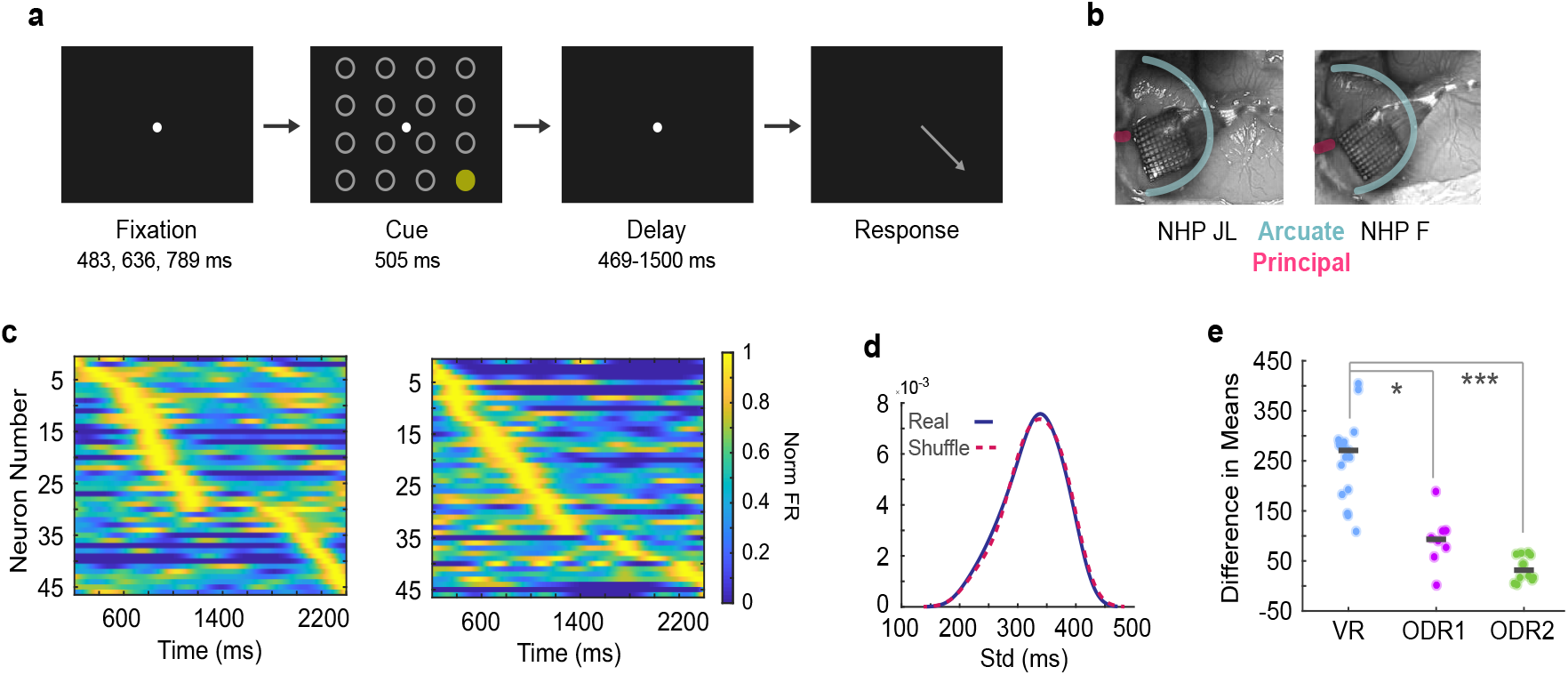
Temporal organization of neural activity during ODR task. **a**, Depiction of ODR task with 16 targets. **b**, Surgical images showing location of Utah arrays implanted in left LPFC of NHP JL and NHP F. **c**, Two trial examples of simultaneously recorded population activity. Normalized firing rate for each neuron is arranged by max firing time. **d**, Real and shuffled distributions of max firing time trial-to-trial deviation for an example session. **e**, Difference in the means between real and shuffled distributions for the virtual reality task and the ODR task. Gray lines indicate median values and dots represent data per session. *p < 0.05, **p < 0.01, ***p < 0.001.

**Fig. 13.**
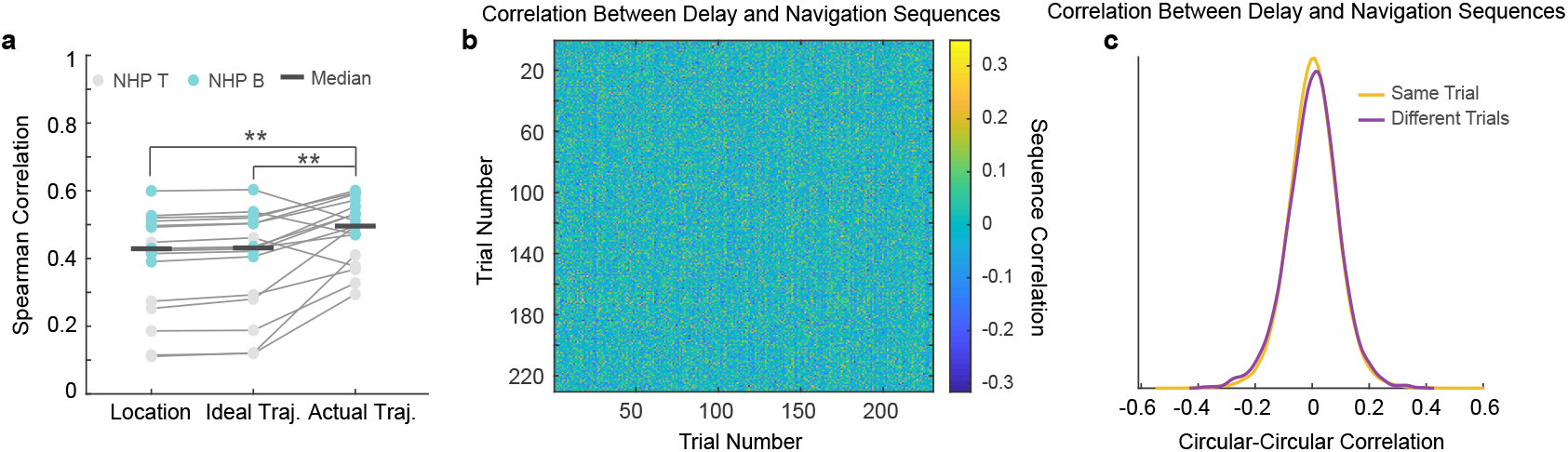
Trajectory analysis. **a**, Correlation between condition centroids and target location, ideal trajectories (i.e., Euclidean distance from start location to targets), or actual trajectories to targets locations. Dots represent data per session. **b**, The correlation between delay and navigation epoch neural sequences between trials for an example session. The diagonal represents sequence correlations within the same trial. If neural sequences repeated between delay and response epochs in the same trial, a clear diagonal of increased correlation values would be present. **c**, Correlation between delay and navigation neural sequences within the same trial and between different trials for all sessions. *p < 0.05, **p < 0.01, ***p < 0.001.

**Fig. 14.**
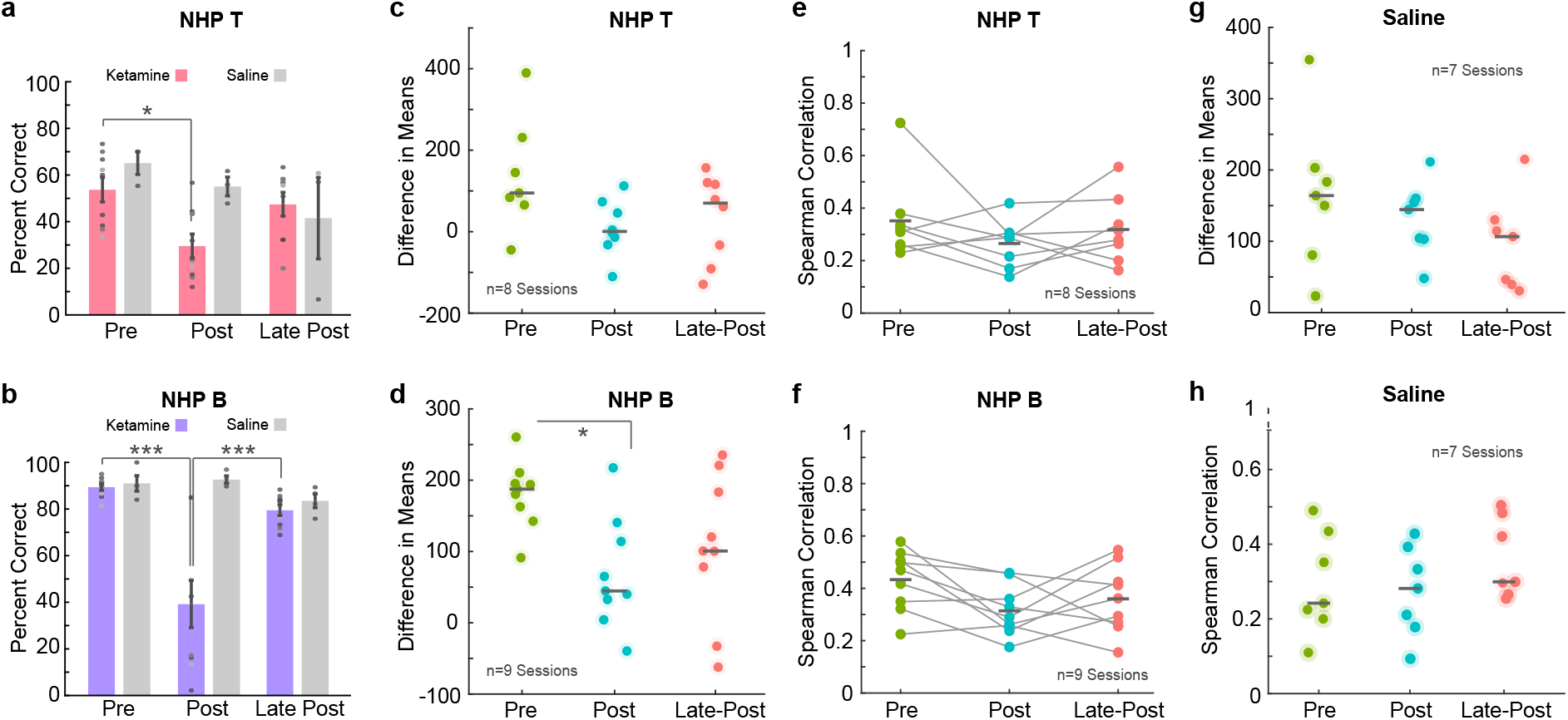
Ketamine and saline control analysis. **a**, Percent of correct trials for ketamine and saline sessions over injection periods for NHP T. Dots represent data for individual sessions and error bars are SEM. **b**, Percent of correct trials for ketamine and saline sessions over injection periods for NHP B. Dots represent data for individual sessions and error bars are SEM. **c**, Difference in the means between real and shuffled distributions of standard deviation values for neuron max firing time between trials. Presented for each ketamine injection period for NHP T. Dots represent data for each session. **d**, Difference in means between real and shuffled distributions for each ketamine injection period for NHP B. Dots represent data for each session. **e**, Correlation between the distance between condition cluster centroids and distance between target trajectories. Correlation values are presented for ketamine injection periods for NHP T. Dots represent data per session. **f**, Correlation values for ketamine injection periods for NHP B. Dots represent data per session. **g**, Difference in mean values between real and shuffled distributions for saline injection periods. Dots represent data for each session (NHP T and NHP B combined). **h**, Correlation between the distance between condition cluster centroids and distance between target trajectories. Correlation values are presented for saline injection periods for both animals combined. *p < 0.05, **p < 0.01, ***p < 0.001.

**Fig. 15.**
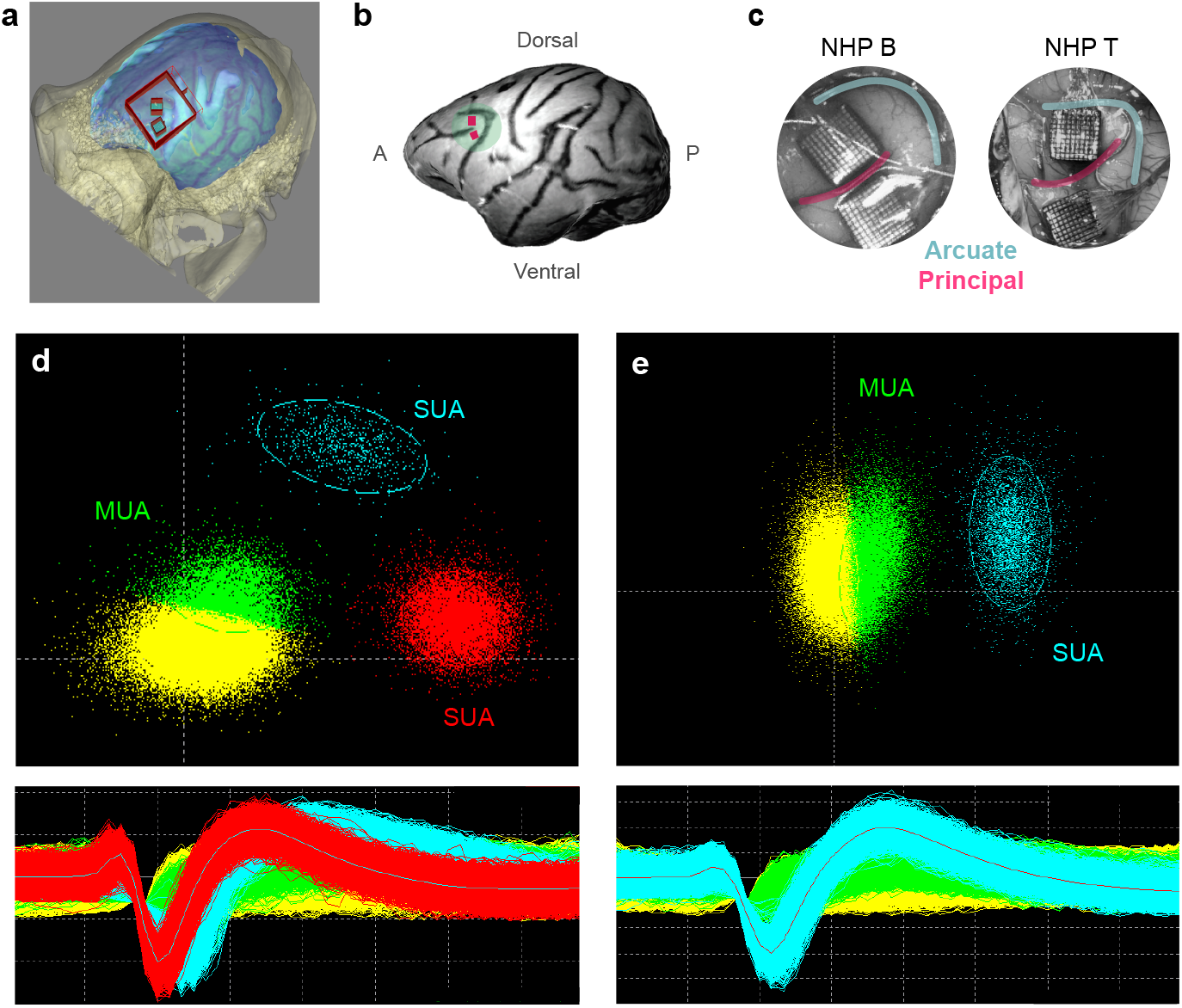
Neural recording setup. **a**, Graphic of presurgical planning procedure showing 3D reconstructed skull and brain based on CT and MRI scans. Electrode array positioning is illustrated in blue with a red outline. The craniotomy is outlined by the larger red box. **b**, 3D modeled brain with electrode array placement in pink. **c**, Surgical images of array implantation in NHP B and NHP T. **d**, Example of spike sorting for one electrode channel. Upper panel represents PCA space and the lower panel represents individual threshold crossing event waveforms. The blue and red clusters represent what we would classify as a single unit. The green cluster would be classified as a multiunit. **e**, Example of spike sorting for one electrode channel. The blue cluster would represent a single unit. The green cluster would represent multiunit activity. This figure is modified from Roussy et al., 2021).

## Methods

We used the same two adult male rhesus macaques (Macaca mulatta) in the main experiment as well as the ketamine and saline experiments (age: 10, 9; weight: 12, 10 kg). The oculomotor delayed response task was recorded from two different male macaques using one multielectrode Utah array implanted in each animal (Leavitt, 2017b, Leavitt, 2018).

### Ethics statement

Animal care and handling (i.e., basic care, animal training, surgical procedures, and experimental injections) were pre-approved by the University of Western Ontario Animal Care Committee. This approval ensures that federal (Canadian Council on Animal Care), provincial (Ontario Animals in Research Act), regulatory bodies (e.g., CIHR/NSERC), and other national standards (CALAM) for the ethical use of animals are followed. The oculomotor delayed response task experiment complied with Canadian policies and regulations and was preapproved by the McGill University Animal Care Committee (Leavitt, 2017b, Leavitt, 2018). Regular assessments for physical and psychological well-being of the animals were conducted by researchers, registered veterinary technicians, and veterinarians.

### Experimental setup

Animals performed the task in an isolated room with no illumination other than the monitor. The room contained no AC power lines and was radiofrequency (RF) shielded. The task was presented on a computer LDC monitor positioned 80 cm from the subjects’ eyes (27” ASUS, VG278H monitor, 1024 × 768 pixel resolution, 75 Hz refresh rate, screen height equals 33.5 cm, screen width equals 45 cm). Eye positions were monitored using a video-oculography system with sampling at 500 Hz (EyeLink 1000, SR Research). Stimulus presentation was controlled through a custom computer program (through Unreal Engine 3). Subjects were seated in a standard enclosed primate chair (Neuronitek) during the experiment and were delivered juice through an electronic reward integration system (Crist Instruments). Prior to the experiments, subjects were implanted with custom fit, PEEK cranial implants which housed the head posts and recording equipment (Neuronitek). See Blonde et al., 2018 for more information. The head posts were attached to the primate chair for head fixation. The experimental setup for the oculomotor delayed response task is outlined in both Leavitt et al. 2017b and Leavitt, 2018.

### Task

The virtual task environment was developed using Unreal Engine 3 development kit, utilizing Kismet sequencing and UnrealScript (UDK, May 2012 release; Epic Games). Details about this platform and the recording setup can be found in Doucet et al., 2016. Movement speed through the environment was fixed.Target locations within the virtual arena were arranged in a 3 × 3 grid and spaced 290 unreal units apart (time between adjacent targets is approximately 0.5 seconds). The perception control variation of the task was identical to the working memory version except that the targets remained onscreen through the trial.

The oculomotor delayed response task was separated into four epochs: fixation, stimulus presentation, delay, and response. The animal began a trial by fixating on a fixation dot and by pressing a lever. The duration of the fixation period was either 482, 636, or 789 milliseconds. A sine-wave grating target then appeared at 1 of 16 randomly selected locations positioned in a 4×4 grid for 505 ms. This was followed by a delay period ranging from 494–1500 ms. The fixation point was removed, cueing the animal to make a saccade to the location of the previously presented target and then to release the lever (see Leavitt et al. 2017b, 2018 for more details).

### Ketamine injection

The ketamine doses were titrated so they did not induce visible behavioral changes in the animals (i.e., nystagmus or somnolence). An intramuscular injection of ketamine (0.25, 0.4, or 0.8 mg/kg) was administered in the hamstring muscles by a registered veterinary technician. Ketamine injections were spaced at least two days apart to allow for washout of the drug. Saline administration was conducted identically with a fixed 0.25 mg/kg dose (See Roussy et al. 2021 for more details).

### Surgical procedure

Custom PEEK implants which housed recording hardware and a headpost were developed and implanted in each animal (Blonde et al., 2018). Brain navigation for surgical planning was conducted using Brainsight (Rogue Research Inc.) (Extended Data Fig. 10a). Two 10×10, microelectrode Utah arrays (96 channels, 1.5 mm in length and separated by at least 0.4 mm) (Blackrock Neurotech) were chronically implanted in each animal. Electrodes were implanted in the left LPFC (anterior to the arcuate sulcus and on either side of the posterior end of the principal sulcus) (Extended Data Fig. 10b, c). Arrays were impacted approximately 1.5 mm into the cortex. Reference wires were placed beneath the dura and a grounding wire was attached between screws in contact with the pedestal and the border of the craniotomy. Surgical procedures were conducted under general anesthesia induced by ketamine and maintained using isoflurane and propofol.

For the oculomotor delayed response task data, a 96-channel Utah array was implanted in each monkey’s left LPFC in the same region that electrodes were implanted for recording during performance of the virtual working memory task (Extended Data Fig. 7b). Detailed surgical methods can be found in Leavitt et al. (2017b, 2018).

### Task performance

Correct trials are trials in which the animal reaches the correct target location within 10 seconds. An incorrect trial occurs if the animal does not reach the target location within 10 seconds. Percent of correct trials is calculated as the number of correct trials divided by the total number of trials. Response time was calculated for correct trials as the navigation start time to the time in which the animal reaches the correct target location.

The optimal trajectory analysis was calculated for correct trials. It is calculated as the real length of the animal’s trajectory to correct target location divided by the optimal trajectory (i.e., the Euclidean distance from the start position to the target location).

For incorrect trials, we calculated the distance from the animal’s final position to the correct target location. Distance values were modified from arbitrary ‘Unreal’ units (the unit system in Unreal Engine Development Kit, Unreal Engine 3, Epic Games) to ‘Unreal’ units divided by the distance between two targets to increase interpretability. A new value of 1 would represent 290 unreal units (the distance between two adjacent targets).

### Eye behavior

Percent of eyes on screen measures the number of eye data points falling on the screen divided by the total number of eye data points. Off screen data points occur when the animal looks off screen or closes their eyes (as occurs during blinking).

Eye data ws classified into fixations and saccades based on a method outlined in Corrigan et al., 2017 that was developed for use in a similar virtual environment. The percent of fixations on target was calculated by the number of fixation events falling within a trial’s target location divided by total number of fixation events. We used a linear classifier (SVM) (Libsvm 3.14, Fan et al., 2008) with 5-fold cross validation to predict target location from eye fixation position data.

The main sequence was calculated by separating saccades into bins of 3° of amplitude, starting at 2° and computing the medians for each bin. The proportion of single units tuned for eye position in both retinocentric and spatiocentric reference frames was calculated using a quadrant binning pattern for a 40°×30° field. A bin had to have at least ten saccades to be acceptable and sessions had at least three acceptable bins.

### Spike processing

Neuronal data was recorded using a Cerebus neuronal Signal Processor (Blackrock Microsystems) via a Cereport adapter. The neuronal signal was digitized (16 bit) at a sample rate of 30 kHz. Spike waveforms were detected online by thresholding at 3.4 standard deviations of the signal. The extracted spikes were semi-automatically resorted with techniques utilizing Plexon Offline Sorter (Plexon Inc.). Sorting results were then manually supervised. Multiunits consisted of threshold-crossing events from multiple neurons with action potential-like morphology that were not isolated well enough to be classified as a well-defined single unit (for spike sorting example see Extended Data Fig. 10d, e). We collected behavioral data across 20 working memory sessions (eight in animal T, twelve in animal B) and neural data across 17 sessions. This yielded a total of 3950 units recorded: 2578 single neurons (346 in animal T, 2232 in animal B) and 1372 multiunits (512 in animal T, 860 in animal B). We collected behavioral data across 18 ketamine-working memory sessions (nine in animal T, nine in animal B) and neuronal data from 17 ketamine-working memory sessions with one session from animal T removed due to incomplete synchronization of neuronal data during the recording. This yielded a total of 2906 units recorded during ketamine-working memory sessions: 1814 single neurons (259 in animal T, 1555 in animal B) and 1092 multiunits (533 in animal T, 559 in animal B).

### Spike density function

Spike density functions (SDFs) were generated by convolving the spike train with a Gaussian kernel (standard deviation=100 ms).

### Time consistent neurons

To qualify time consistent neurons, we created SDFs combined between electrodes arrays over the entire trial time using neurons with firing rates above 0.5 Hz. SDFs were created for each condition that contained at least five trials. We calculated the peak firing time for each neuron in the population, calculated the standard deviation of the peak firing time for each neuron over all trials in a condition and created a probability distribution from the standard deviation values. We shuffled the peak firing times for each neuron from trial to trial so that the peak firing time no longer aligned for any one neuron. We created a shuffled probability distribution. We calculated the difference in mean values between the real and shuffled distributions to get the mean difference value. To calculate the standard deviation values plotted in Extended Data 4a, we calculated trial-trial standard deviation of peak spike time for the target condition in which each neuron fired the most consistently during correct trials (i.e., lowest deviation). The same conditions were used for shuffled data and for incorrect trials.

ODR1: This data was collected from the same animals and electrodes as our naturalistic VR task. This task contained 16 targets (Extended Data Fig. 7a) and was a variation of a traditional ODR task in which the fixation point changes location across trials, resulting in many different task conditions with varying combinations of fixation location and target location. For this reason, we grouped trials with target locations within the same quadrant (same direction saccade). To match the task structure of the VR task, we did not use data from the fixation period.

ODR2: This data was collected from NHPs JL and F (Extended Data Fig. 7b) using one Utah array implanted in the left LPFC (same region as NHP B and T). This task contained 16 targets with a consistent central fixation point (Extended Data Fig. 7a). To match the task structure of the VR task, we did not use data from the fixation period. Since the ODR2 task had jittered delay epoch timing, we used trials with delay periods > 1000 ms and included the first 1000 ms of the epoch.

### Sequence representation

Each trial was represented as a complex-valued vector by mapping the time of maximum spike density of each neuron to a phase value between - and. For the main analysis, only cells with peak firing time during the delay (working memory) period were included in the sequences. To compare across trial epochs, cue and navigation sequences were also considered, including only the cells with peak firing time in the cue and navigation periods respectively.

### Dimensionality reduction

For each recording session, a correlation matrix was created by computing the correlation coefficient between each pair of phase vectors representing single trial sequences. The eigenspectrum of this matrix was computed, and the eigenvalues sorted in descending order of modulus. The correlation matrix was then projected onto the eigenvectors corresponding to the first three sorted eigenvalues to generate a low-dimensional summary of the data in 3D-space. The points in this projection each correspond to one trial, and their positions are determined by the relative similarity of the corresponding sequences. The centroids of the clusters corresponding to each trial condition were then determined, and the matrix of Euclidean distances between the centroids was computed.

The points corresponding to a specific target location l defined a cluster, and the centroids were computed, resulting in one coordinate triple, *c_l_* = (*x_i_,y_i_, z_i_*)*ϵR*^3^, corresponding to each target location. The matrix, D, of Euclidean distances between each pair of centroids was then computed, with *D_i,j_* = *dist*(*c_i_, c_j_*) = √((*x_j_* – *χ_i_*) – (*y_j_* – *y_i_*) + – *z_i_*)^2^).

In this way, the spiking data from each recording session was reduced to a 9×9 distance matrix.

### Correlation analysis

For each recording session, mean trajectories followed by the subject to each target location were obtained by averaging the group of correct trajectories to that target (excluding outlier trajectories with zscore > 1 of mean Frechet distance to other trajectories in the group) (Alt & Godau, 1995). A 9×9 distance matrix was then created from the Frechet distances between each of the mean trajectories. In this way, each recording session was described by two 9×9 normalized distance matrices, one representing the relationships between target centroids in 3-space, and the other representing distances between behavioral trajectories to the target locations in the virtual environment. The correlation between the two distance matrices was then computed, which measures the similarity between the neuronal representations of the targets and the physical trajectories to them.

### Target columns

Since centroids of targets in the same column tended to cluster together, some subsequent analyses group sequences by target column rather than trial condition. In these analyses, the clusters corresponding to targets in the same column were combined to produce 3 centroids instead of 9. Furthermore, due to this structure observed in the data, a null model for comparison was created by shuffling the target columns while preserving the target rows, thereby destroying the correlation while preserving more of the structure of the data.

### Dimensionality reduction for ODR

When repeating the analysis for the oculomotor delayed response task, 16 targets were included so the data was reduced to a 16×16 distance matrix describing the relationships between the neuronal representations of the target locations. Furthermore, since the ODR task does not contain a navigation component, the distance matrix of the projection centroids was compared to the matrix of Euclidean distances between targets (rather than a Frechet distance matrix).

### Projection classification analysis

We used a simple classifier based on our computational approach with 5-fold crossvalidation to classify sequences on a single trial basis. The classifier assigns labels to points in the projected 3-space, assigning each trial in the test set (20% of trials) to the centroid of trials in the training set (80% of trials) which has the minimum Euclidean distance to the trial in question. In the supervised version, the training set centroids are determined using the trial condition labels. The unsupervised version uses K-means clustering to determine the training set centroids.

The same method is used for all classifiers. To classify single trials by trial condition (9 targets), the supervised classifier was necessary. To classify single trials by target column (3 columns), the unsupervised version was used. To classify single sequences by trial epoch (3 epochs), the cue, delay, and response sequences (see ‘Sequence Representation’) were all used to create the correlation matrix (i.e., the correlation between each pair of sequences from all 3 epochs was used to generate the projection), and the centroids are defined by sequence epoch clusters rather than trial condition clusters.

### Trajectory analysis

We repeated the centroids analysis described above, but replaced the distance matrix describing mean trajectories with two other task-relevant measures. First, we constructed geometrically ‘ideal’ trajectories straight from the start location to each target, and repeated the analysis using Frechet distance between trajectories. Second, we used the matrix of Euclidean distances between target locations. Unlike the mean trajectory analysis, these measures only described the task set up and did not include behavioral data.

To determine whether the sequences represented motor planning during the delay that was replayed during navigation, we compared the sequences specific to the delay epoch with those from the navigation epoch. For each recording session, we computed the correlation between each delay-navigation sequence pair. We then considered the pairs in which the delay sequence and navigation sequence corresponded to the same trial separately from the pairs where the two sequences came from different trials. To consider this measure across recording sessions, we computed the distributions of correlation values for each group (same trial versus different trial).

### Single versus multiple contribution

In the main analysis, we assume each cell contributes only once to the sequence representation, at its time of peak firing. As such, each cell participates in only one of the cue, delay, or navigation epoch sequence. To test the validity of this assumption, we repeated the analyses for sequences in which each cell participated in all three epochs, by considering the time of max firing within each epoch.

### Removing cells

For percentages increasing from 10% to 90% in 10% increments, we removed a percentage of cells contributing to the delay sequence in each trial. The correlation analysis was then performed for the sequences with removed cells. This process was repeated for 10 iterations of both correct trials and incorrect trials, and correlation values were averaged across iterations to produce the plot. The difference between results for correct and incorrect trials is also plotted.

